# Maternal low-fat and high-fat diet decreases survival and alters cytokine signaling in neonatal mice with *Staphylococcus epidermidis* sepsis

**DOI:** 10.1101/2025.07.30.667518

**Authors:** Lauren Bodilly, Sarah Weiner, Jennifer Bermick

## Abstract

**Objective:** Maternal malnutrition increases susceptibility to sepsis and mortality in neonates. The reason for this increased susceptibility remains unknown. We aimed to evaluate bacterial burden and serum cytokine levels in septic neonatal mice born to dams with malnutrition.

**Methods:** 6-week-old C57BL/6 dams were placed on a low-fat (LFD) (10% kcal from fat), control (CD) (18% kcal from fat), or high-fat (HFD) (60% kcal from fat) diet for 3 weeks prior to breeding. Sepsis was induced in P4-P6 offspring via intraperitoneal *Staphylococcus epidermidis* injection. Mice were monitored for survival. At 12h after sepsis, serum and peritoneal wash fluid were collected for bacterial count and serum cytokine levels. In the absence of infection, P4-P6 offspring had untargeted serum metabolomics performed.

**Results:** Septic offspring of dams fed LFD and HFD had significantly higher mortality than offspring of dams fed CD. There was no difference in serum or peritoneal wash bacterial loads. Maternal diet and *Staphylococcus epidermidis* sepsis caused changes in basal serum cytokine levels, with HFD causing decreased cytokine elevation during sepsis. Maternal LFD and HFD altered similar metabolomic pathways in offspring.

**Conclusion:** Maternal LFD and HFD decrease survival during neonatal sepsis and alter serum cytokines and the metabolome, supporting a role for maternal nutrition in neonatal immune function and infection susceptibility.

## 1. Introduction

Sepsis is a leading cause of morbidity and mortality in neonates.(1) Sepsis is a syndrome in which a life-threatening infection leads to a dysregulated host immune response.(2) Neonatal sepsis can be early-onset, within the first 72 hours of life, or late-onset, occurring after 72 hours of life. Coagulase-negative staphylococcus, including *Staphylococcus epidermidis* (*S. epidermidis*), is a common pathogen causing late-onset sepsis. Neonatal risk factors for sepsis include, but are not limited to, prematurity, low birth weight, and meconium-stained amniotic fluid.(3) Hospital-associated risk factors include presence of a central venous line and mechanical ventilation. Additionally, there are maternal risk factors such as premature rupture of membranes, infection, and maternal weight.(3-6)

Maternal malnutrition is a global epidemic that can be categorized as undernutrition, overnutrition, or micronutrient deficiency or excess. Currently 9.1% of women suffer from undernutrition and 32.5% of women suffer from overnutrition, with rates of overnutrition continuing to rise globally.(7) Neonates born to mothers with undernutrition or overnutrition have increased rates of pre-term birth, growth restriction or excess, sepsis, and mortality.(4, 8, 9) However, there remains a significant knowledge gap in understanding why maternal malnutrition increases the risk of sepsis in neonates.

The fetal programming hypothesis suggests that alterations in fetal nutrition, related to maternal nutrition and other exogenous factors, determine an individual’s long-term immunologic and metabolic health.(10) There is evidence that umbilical cord blood cell lineages are different in infants born small-for-gestational age compared to those born appropriate-for-gestational age; however, data was not collected on maternal nutrition or weight.(11) A murine maternal undernutrition model demonstrated that offspring CD4+ T cells skewed toward the TH2 lineage and resulted in an asthma-like phenotype, highlighting the immune effects of maternal undernutrition.(12)

Several studies on maternal overnutrition demonstrate altered placental immunity.(13-15) Additionally, umbilical cord blood monocytes and myeloid dendritic cells from obese pregnancies have decreased lipopolysaccharide (LPS) induced responses.(16, 17) Dampened monocyte responses were also shown following *Escherichia coli* (*E.coli*) stimulation and were replicated in fetal peripheral monocytes and tissue-resident macrophages in a rhesus macaque model of maternal obesity.(18) In a murine model of maternal overnutrition, offspring born to dams on obesogenic diets had increased susceptibility to bacterial infection, although this infectious challenge occurred after weaning rather than during the neonatal period.(19)

Given the paucity of information on the impact of maternal malnutrition on neonatal immunity, we implemented a murine model of maternal malnutrition by exposing dams to low-fat, normal-fat, or high-fat diet and infected their offspring with *S. epidermidis* to induce late-onset sepsis. We hypothesized that neonates born to low-fat and high-fat diet fed dams would have dampened immune responses and decreased survival during *S. epidermidis* sepsis.

## 2. Methods

### 2.1 Animals

The investigations were conducted in accordance with the ARRIVE guidelines and were approved by the Institutional Animal Care and Usage Committees at the University of Iowa.(20) Mice were housed in a University of Iowa vivarium. Food and water were provided ad libitum. Female and male C57BL/6 mice aged six weeks were obtained from The Jackson Laboratories (Bar Harbor, ME, USA) or were offspring of in-house breeding pairs of C57BL/6 dams and sires. Neonatal male and female mice born in-house were used for all experiments.

### 2.2 Model of maternal malnutrition

Six-week-old C57BL/6 female mice were selected by cage to be placed on a low-fat diet (LFD; Research Diets Inc D12450J; 10% kcal provided by fat), on control-fat diet (CD; Research Diets Inc D17030603; 18% kcal provided by fat), or high-fat diet (HFD; Research Diets Inc D12492, 60% kcal provided by fat) (Table 1). The control-fat diet is a purified diet that is matched in kcal provided by fat to the grain control chow, 7913 NIH-31 Irradiated Modified Open Formula Mouse Diet, provided by our Office of Animal Resource and matched in protein and micronutrients to the low and high fat diets. Body weights were monitored weekly for 3 weeks and the females were then mated with C57BL/6 males. After breeding, females were independently housed and assessed for pregnancy by intravaginal plug formation and visual inspection. Female mice remained on their designated diet for the entire experiment. Male mice were exposed to the LFD, CD, or HFD during the week of mating. No blinding occurred during the study as the diets are different colors and easy to distinguish.

**Table 1.** Diet composition of D12450J (LFD), D17030603 (CD), and D12492 (HFD) from Research Diets, Inc.

| Diet | D12450J | D17030603 | D12492 |
| --- | --- | --- | --- |
| %kcal |  |  |  |
| Protein | 20 | 20 | 20 |
| Fat | 10 | 18 | 60 |
| Carbohydrate | 70 | 62 | 20 |
| Kcal/gram | 3.85 | 4.0 | 5.2 |
| Ingredients (gram) |  |  |  |
| Casein, 30 Mesh | 200 | 200 | 200 |
| DL-Methionine | 3 | 3 | 3 |
| Corn Starch | 506.2 | 427.5 | 0 |
| Maltodextrin 10 | 125 | 125 | 125 |
| Sucrose | 68.8 | 68.8 | 68.8 |
| Cellulose, BW200 | 50 | 50 | 50 |
| Soybean Oil | 25 | 25 | 25 |
| Lard | 20 | 55 | 245 |
| Mineral Mix S100026 | 10 | 10 | 10 |
| Calcium Carbonate | 13 | 13 | 13 |
| Sodium Chloride | 5.5 | 5.5 | 5.5 |
| Potassium Citrate, 1 H2O | 16.5 | 16.5 | 16.5 |
| Vitamin Mix V10001 | 10 | 10 | 10 |
| Choline Bitartrate | 2 | 2 | 2 |

### 2.3 Body Composition

Total body, fat, lean, and fluid mass were quantified by NMR spectroscopy using a Bruker mini-spec LF 90II instrument (Bruker Corporation, Billerica, MA). To analyze body composition, mice were placed into a restraint tube and inserted into the NMR machine.

### 2.4 Staphylococcus epidermidis

*S. epidermidis* (Schroeter) Migula (ATCC 12228) was cultured at 37 °C overnight in nutrient broth as seed culture to inoculate a fresh broth that was grown for 2 hours. Spectrophotometry was used to estimate bacterial concentration of broth, and a growth curve was used to determine amount of broth needed to be pelleted to achieve approximately 2×10^9^ cfu/mL. Pellet was suspended in 1mL of PBS (no Ca^2+^/no Mg^2+^) to produce inocula. Inocula was plated in 1:10 dilutions and incubated for 24 hours. Serial dilutions resulted in a final density of 2×10^7^-1×10^9^ cfu/mL which corresponded to a dose of 1×10^7^-5×10^8^ cfu/injection.

### 2.5 Neonatal Sepsis

Neonatal mice were chosen to be infected or not infected based on litter due to the need to remain with their mothers. P4-P6 neonatal mice were weighed. Mice less than 2.25g were excluded from the study. P4-P6 neonatal mice infected with *S. epidermidis* solution via intraperitoneal (i.p.) injection (50uL/mouse). They remained with their mother and littermates following infection.

### 2.6 Survival Studies

After injection of *S. epidermidis*, mice were monitored for survival for 5 days. Mice were monitored every 6 hours for the first 24 hours and then every 12 hours thereafter. An injury severity scale was used to assess the severity of symptoms. Mice were graded on a scale of normal, decreased, or absent for presence of milk spot, vigor, and if located near littermates. If they scores were absent or significantly decreased across all three categories, then they were compassionately euthanized.

### 2.7 Blood and Tissue Harvest

Neonates were euthanized at 12 hours post infection for blood and tissue harvest. Neonates were decapitated and whole blood collected with 50uL of heparin in a 24 well plate. Whole blood samples were centrifuged at 3000 g for 15 min at room temperature. The serum was aliquoted into separate microcentrifuge tubes and samples were frozen at −80 °C until analysis.

Dams were given a sublethal dose of sodium pentobarbital (150 mg/kg i.p.) and intracardiac puncture was performed to obtain whole blood once the mouse was sufficiently anesthetized. As described above, whole blood samples were centrifuged, and the serum was frozen at −80 °C. Dams were cervically dislocated after intracardiac puncture.

### 2.8 Bacterial Count

Following decapitation, whole blood was collected as described above. To collect peritoneal fluid, 150uL of sterile PBS was injected into the peritoneal cavity of neonatal mice and removed twice per animal. All samples were serially diluted in sterile PBS. The following dilutions were plated on nutrient agar plates: 1:10 and 1:100 and peritoneal fluid 1:10 through 1:1,000,000. Colony-forming units were counted after 24h.

### 2.9 Cytokines

Cytokine levels were quantified in neonatal serum samples using bead-based multiplexed technology. Protein levels of the cytokines G-CSF, GM-CSF, TNF-α, IL-1β, IL-6, IL-10, MIP-1α, MIP-1β, MCP-1, and RANTES were quantified using a 23-cytokine multiplexed Bio-Plex assay (#M60009RDPD) on a Bio-Plex 200 machine following the manufacturer’s instructions (Bio-Rad, Hercules, CA, USA). Data is not shown for the rest of the panel.

### 2.10 Metabolomic Profiling

All metabolite profiling was performed by Metabolon, Inc (Durham, NC). All serum was collected from P4-P6 mice and stored at -80 until sample processing. Briefly, serum was thawed and pooled from 3-6 mice per dietary intervention group to achieve 150uL/sample. Samples were sent to Metabolon, Inc for non-targeted metabolic profiling using UHPLC-MS/MS. A total of 1,034 different metabolites were identified by automated comparison of the serum samples to an in-house library of chemical standards.

### 2.11 Metabolic Pathway Analysis

Metabolomic data was analyzed using Metaboanalyst v6.0. To perform statistical analysis (one factor) the original data file from Metabolon was saved and edited to include only offspring of LFD-fed and grain CD-fed dams or HFD-fed and grain CD-fed dams. Data was median-centered, log-transformed, and pareto-scaled. Principal Component Analysis was performed with statistical significance evaluated using PERMANOVA and distributions computed using the Euclidean distance from the center. Metabolic pathway analysis was conducted to better understand the functional impact of maternal diet on neonatal metabolism. This feature combines pathway enrichment analysis and pathway topology analysis as previously described.(21) This was performed using the same data files and data pre-processing strategies as described above. The analysis was performed using the following parameters: (1) enrichment analysis was performed using the global test, (2) centrality was measured using Relative Betweenness, and (3) Mus musculus (house mouse) the Kyoto Encyclopedia of Genes and Genomes (KEGG) database were used as a reference for metabolic pathways.

### 2.12 Statistical Analysis

Data were analyzed using GraphPad Prism. For data comparison among only two groups, statistical analysis was performed using *t*-test for parametric data and Mann-Whitney U for non-parametric data. For data comparison among three or more groups, statistical analysis was performed using two-way ANOVA with the Holm-Sidak multiple comparisons. If data were not normally distributed, they were log-transformed and analyzed for normality prior to analysis. Data are expressed as mean±SD for parametric data and median (IQR) for nonparametric data in the text and figures. Survival analysis was performed by log rank test. A value of *P* ≤ 0.05 was considered significant.

## 3. Results

### 3.1 Maternal malnutrition alters maternal and/or neonatal weight and body composition

To model maternal undernutrition and overnutrition, dams were maintained on low-fat (LFD), control-fat (CD), or high-fat (HFD) diets throughout gestation and lactation. After 3 weeks on diet, HFD-fed dams exhibited significantly increased body weight and altered body composition, including elevated fat mass and reduced lean mass percentage, relative to CD-fed controls (Figure 1A–B). In contrast, LFD-fed dams did not differ significantly in weight or body composition compared to CD-fed dams.

**Figure 1.**
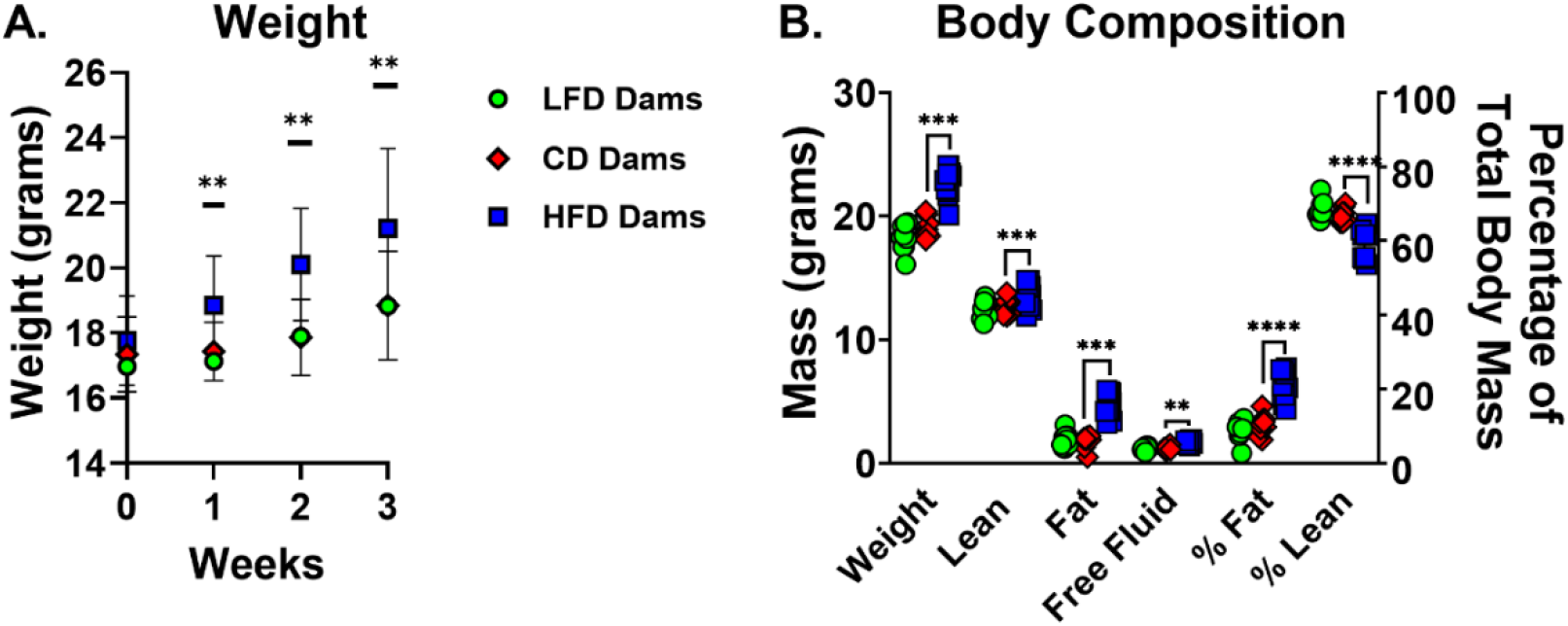
Maternal weight and body composition is impacted by high-fat diet but not low-fat diet. 6-week-old C57BL/6 dams were placed on a low-fat diet (LFD) (10% kcal from fat), control-fat diet (CD) (18% kcal from fat), or high-fat diet (HFD) (60% kcal from fat) for 3 weeks prior to breeding. (A) LFD fed dams weigh the same as CD fed dams. HFD fed dams weighed more than CD fed dams from week 1 through week 3 (n=15/group). Data presented as mean with SD bars. (B) After 3 weeks of dietary intervention, separate cohorts of dams underwent NMR to assess body composition (n=10/group). There was no difference in body composition in LFD and CD fed dams. HFD fed dams had significantly higher weight, lean mass, fat mass, and total body water mass compared to CD fed dams. They also had higher fat mass percentage and lower lean mass percentage. \*\**P*<0.01, \*\*\**P*<0.001, \*\*\*\**P*<0.0001. Data were analyzed using t-tests comparing LFD to CD and HFD to CD.

There was no significant difference in litter size on P0 and survival to P4-6 (day of sepsis) (Figure 2A). Despite comparable maternal body composition, LFD-fed dams produced significantly smaller offspring at the time of sepsis induction compared to CD-fed dams (Figure 2B). HFD-fed dams did not significantly differ in litter or offspring size compared to the other groups. Despite the lack of statistical significance, HFD-fed dams trended toward heavier offspring. These data suggest that maternal diet macronutrient composition, rather than caloric intake, may underlie the observed offspring phenotype.

**Figure 2.**
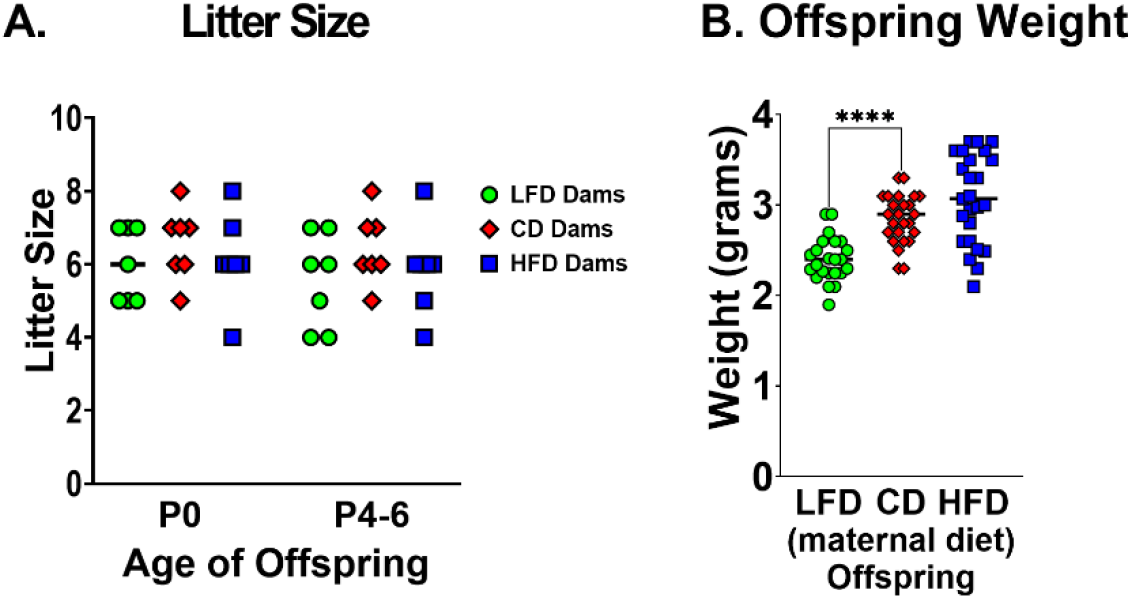
LFD fed dams had significantly smaller litter sizes and lighter offspring at time of sepsis. After 3 weeks of dietary intervention, C57BL/6 dams were bred with C57BL/6 males. Dams remained on their prescribed diet throughout gestation and lactation. Males were exposed only during the time of breeding and otherwise received standard chow. Offspring were counted and weighed at time of sepsis (P4-P6). (A) LFD fed dams had significantly smaller litter sizes compared to CD fed dams. There was no difference in HFD fed and CD fed dam litter sizes (n=9-10 litters/group). (B) Offspring born to dams fed a LFD weighed significantly less than offspring born to dams fed a CD. Although it is not statistically significant, offspring of HFD-fed dams trends toward weighing more than offspring of CD-fed dams (n=23-25/group). Bars represent median. Individual data points presented. \**P*<0.05, \*\**P*<0.01, \*\*\**P*<0.001, \*\*\*\**P*<0.0001. Data were analyzed using t-tests comparing low-fat diet to control-fat diet and high-fat diet to control-fat diet.

### 3.2 Maternal malnutrition decreases neonatal survival during *S. epidermidis* sepsis independent of bacterial burden

To assess the impact of maternal diet on neonatal susceptibility to infection, offspring were challenged with intraperitoneal *S. epidermidis* and monitored for survival. First, we compared survival of offspring of dams fed the standard chow to offspring of dams fed the CD, and there was no significant difference in survival (Figure 3A). Offspring of both LFD-and HFD-fed dams exhibited significantly reduced survival compared to offspring of CD-fed dams (Figure 3B). The greatest decrease in survival is seen between 12- and 24-hours post-infection. These findings suggest that both maternal under- and overnutrition compromise neonatal host defense mechanisms.

**Figure 3.**
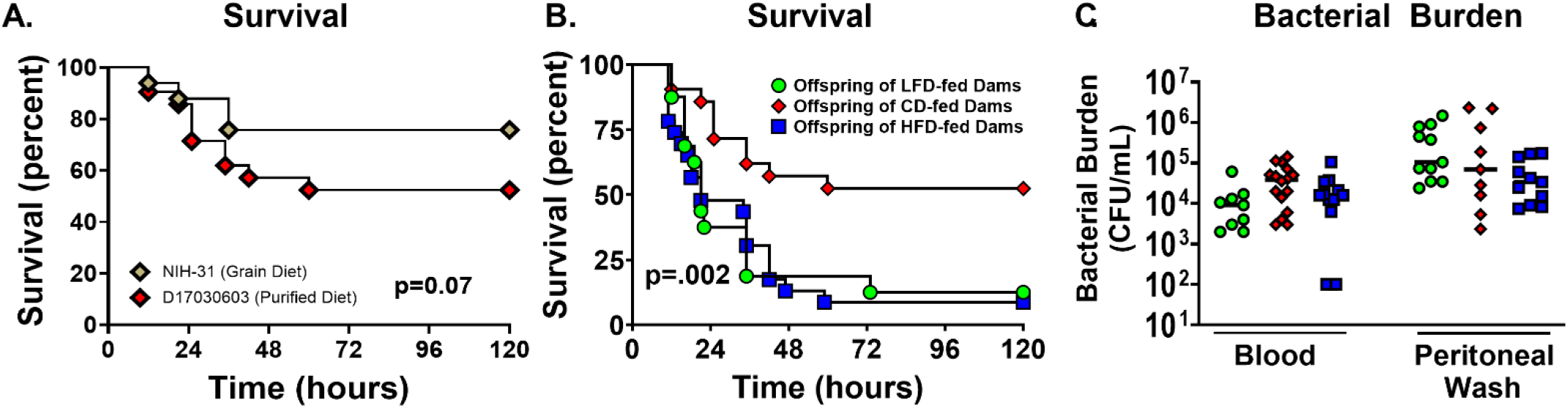
Offspring born to LFD and HFD fed dams have significantly lower survival with no difference in bacterial burden. Sepsis was induced in P4-P6 offspring with i.p. *S. epidermidis*. Offspring were monitored for 5 days for survival. (A) Survival was not different in offspring of mice born to dams fed the standard grain diet (NIH-31) or the purified control diet (D17030603) (n=21-33/group). (B) Survival was significantly lower in offspring of LFD fed dams (13%) and HFD fed dams (9%) vs offspring of CD fed dams (52%) (n=16-23/ group). \*\**P*<0.01 using the log-rank test. Results are combined from 3 experiments. (C) There was no difference in bacterial count in blood or peritoneal fluid (n=12-24/group). Results are combined from 2 experiments.

To determine whether decreased survival was attributable to impaired bacterial clearance, bacterial load was quantified in blood and peritoneal fluid 12 hours after *S. epidermidis* injection. No significant differences in colony-forming units were observed across dietary groups in blood or peritoneal fluid (Figure 3C).

### 3.3 Maternal diet and *S. epidermidis sepsis* modulate cytokine profiles in neonatal serum

We hypothesized that maternal malnutrition causes a dampened immune response in neonates. In the absence of infection, offspring of LFD- and HFD-fed dams exhibited significantly reduced serum levels of key cytokines involved in innate immunity, including G-CSF, GM-CSF, IL-6, and IL-10, compared to CD-fed dam offspring (Figure 4A). Following infection, all neonatal groups mounted significant increases in serum cytokines and chemokines, including G-CSF, GM-CSF, TNF-α, IL-6, MIP-1α, MIP-1β, and RANTES (Figure 4B). This confirms the activation of a systemic inflammatory response to *S. epidermidis* across maternal dietary conditions.

**Figure 4.**
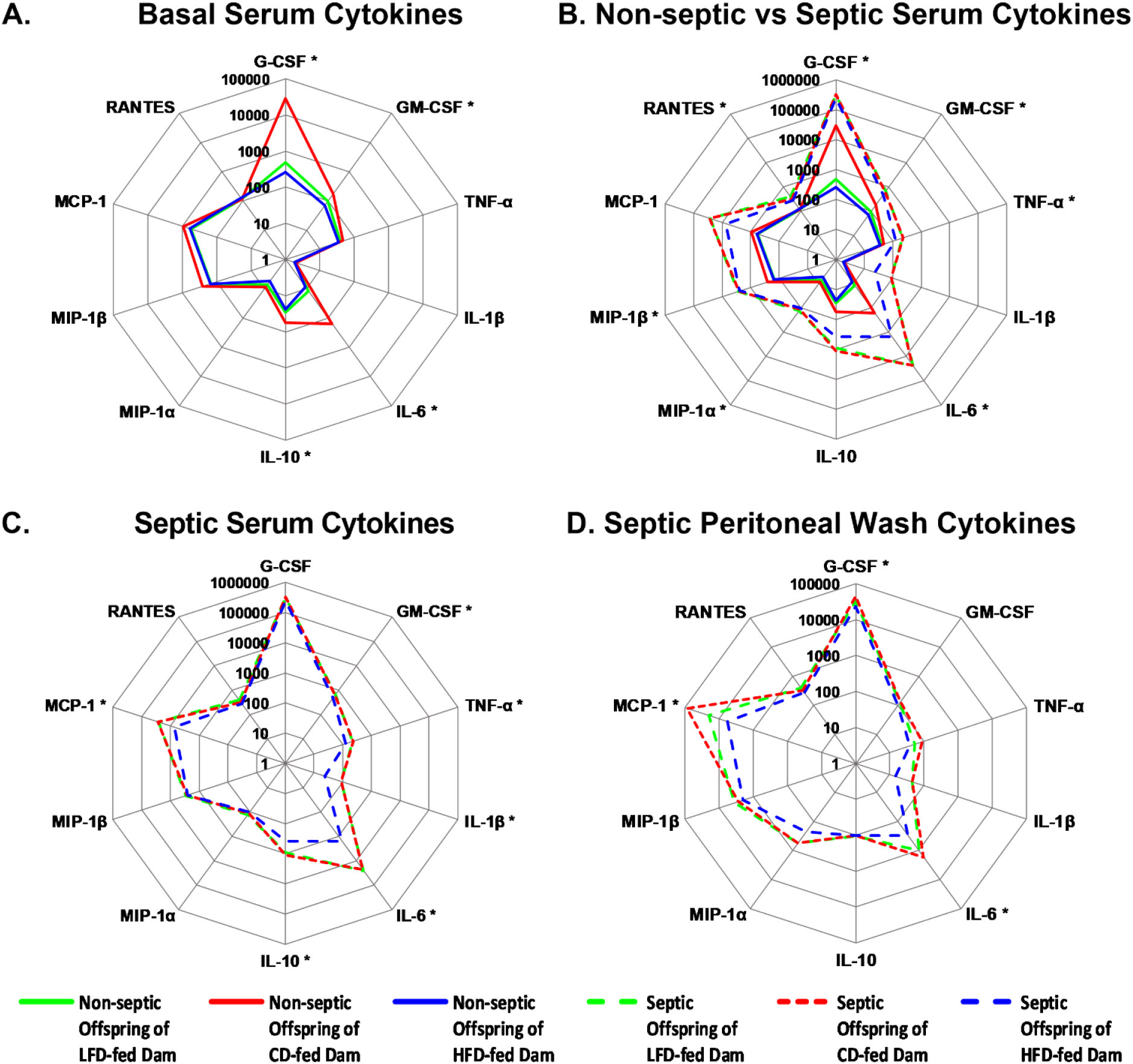
Maternal diet and *S. epidermidis sepsis* influence serum cytokine levels. (A) In the absence of infection, offspring born to dams fed a LFD and HFD have significantly lower levels of G-CSF, GM-CSF, IL-6, and IL-10 compared to offspring born to dams fed a CD (n=8/group; offspring from 1-2 litters). Data were analyzed using t-tests to compare non-septic offspring of LFD dams or HFD dams to offspring of CD dams. (B) Offspring of LFD, CD, and HFD-fed dams have significantly higher levels of serum cytokines 12 hours after *S. epidermidis* sepsis (n=8/group; offspring from 2 litters). Data were analyzed using Two-way ANOVA to compare non-septic and septic offspring of LFD dams or HFD dams to offspring of CD. (C) Septic offspring of LFD fed dams do not have significantly different serum cytokine levels compared to septic offspring of CD fed dams at 12 hours after sepsis. Septic offspring of HFD fed dams have significantly lower levels of key cytokines that regulate innate immune response compared to septic offspring of CD fed dams at 12 hours after sepsis. (D) Septic offspring of LFD fed dams do not have significantly different peritoneal wash cytokine levels compared to septic offspring of CD fed dams at 12 hours after sepsis. Septic offspring of HFD fed dams have significantly lower levels of G-CSF, IL-6, and MCP-1 compared to septic offspring of CD fed dams at 12 hours after sepsis. Data were analyzed using One-way ANOVA to compare non-septic and septic offspring of LFD dams or HFD dams to offspring of CD dams. Data are presented on a radar chart. Solid lines represent median of non-septic mice and dashed lines represent median of septic mice. \**P*<0.05, Green lines= offspring of LFD dams; red lines= offspring of CD dams; blue lines= offspring of HFD dams.

Despite similar bacterial loads, septic offspring of HFD-fed dams exhibited blunted cytokine responses compared to CD-fed dams. At 12 hours after *S. epidermidis* infection, offspring of HFD-fed dams had significantly reduced levels of serum GM-CSF, TNF-α, IL-1β, IL-6, IL-10, and MCP-1 compared to offspring of CD-fed dams (Figure 4C). Similarly, septic offspring of HFD-fed dams had significantly reduced levels of peritoneal wash G-CSF, IL-6, and MCP-1 compared to septic offspring of CD-fed dams (Figure 4D). These data suggest that maternal overnutrition impairs neonatal cytokine signaling during infection, potentially contributing to decreased survival.

In contrast, septic offspring of LFD-fed dams demonstrated cytokine and chemokine profiles comparable to offspring of CD-fed dams at 12 hours after *S. epidermidis* infection. Septic offspring had robust elevations in IL-1β, IL-6, IL-10, and MCP-1 (Figure 4C). There was no difference in peritoneal wash cytokine levels in septic offspring of LFD and CD-fed dams (Figure 4D). These findings indicate that while maternal LFD alters several basal serum cytokine levels, it does not significantly impair the acute cytokine response to infection.

### 3.4 Maternal diet reprograms the neonatal serum metabolome

Given the lack of a clear mechanism for decreased survival in septic pups born to malnourished mothers, we sought to determine if changes in the neonatal metabolome could be playing a role. Untargeted metabolomics was performed on serum from offspring without sepsis to evaluate for baseline differences. Between-class principal-component analysis with k-means statistical comparison revealed there was a significant difference in the composition of the serum metabolome in offspring born to GD-fed dams and LFD-fed dams (Figure 5A, p=0.007). [PERMANOVA] F-value: 6.2201; R-squared: 0.4705; p-value (based on 999 permutations): 0.007.

**Figure 5.**
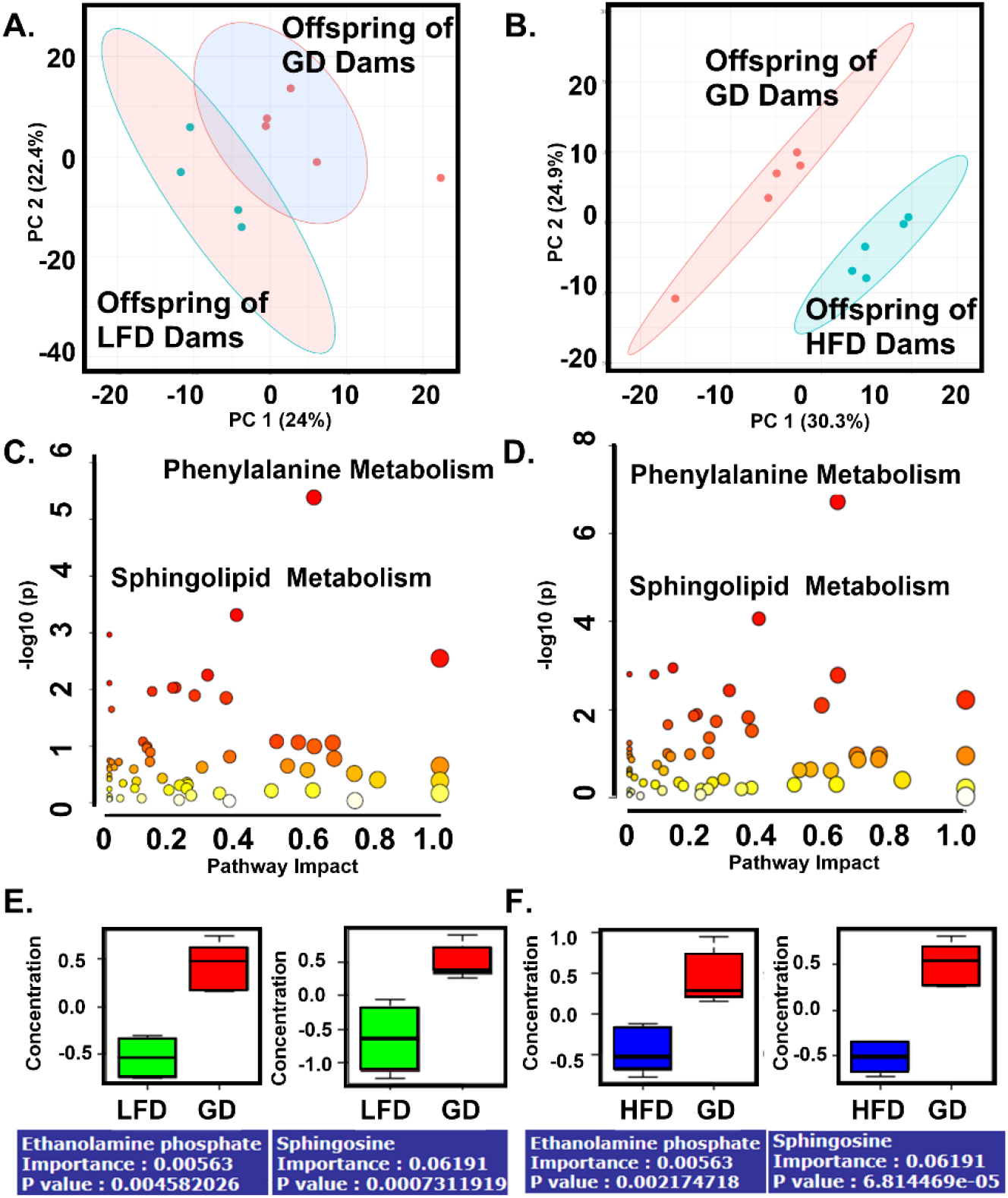
Offspring born to dams fed a LFD or HFD have significantly different serum metabolomes compared to offspring born to dams fed a GD. Serum from non-septic P4-P6 offspring was pooled (3-6 offspring per sample) and analyzed using UHPLC-MS/MS performed by Metabolon, Inc. (A) Principal-Component Analysis (PCA) with k-means statistical comparison demonstrates a significant difference in serum metabolome in offspring of LFD and GD-fed dams. (B) Principal-Component Analysis (PCA) with k-means statistical comparison demonstrates a significant difference in serum metabolome in offspring of HFD and GD-fed dams. (C) Pathway Impact Analysis of offspring of LFD-fed dams. (D) Pathway Impact Analysis of offspring of HFD-fed dams. (E) Sphingolipid metabolites are lower in offspring of LFD-fed dams. (F) Sphingolipid metabolites are lower in offspring of HFD-fed dams.

There was also a significant difference in the composition of the serum metabolome in offspring born to GD-fed dams and HFD-fed dams (Figure 5B, p=0.012). [PERMANOVA] F-value: 8.7225; R-squared: 0.5216; p-value (based on 999 permutations): 0.012. Pathway enrichment analysis identified phenylalanine and sphingolipid metabolism as significantly impacted in both malnourished groups, suggesting shared metabolic disruptions that may underlie altered immune phenotypes (Figures 5C and 5D). There were multiple metabolites in the sphingolipid metabolism pathway that were altered in the same direction in offspring of LFD and HFD-fed dams (Figures 6E and 6F).

## 4. Discussion

This study demonstrates that both maternal undernutrition and overnutrition, modeled through low-fat and high-fat diets respectively, significantly impair neonatal immune responses and increase susceptibility to sepsis.

Despite divergent effects on maternal body composition, both maternal LFD and HFD resulted in decreased neonatal survival following *S. epidermidis* infection, independent of bacterial burden. These findings suggest that maternal diet exerts a profound influence on neonatal immune programming, likely through mechanisms beyond simple nutrient availability or pathogen load.

This study is the first, to our knowledge, to examine the impact of maternal dietary fat composition on neonatal sepsis outcomes. Maternal malnutrition—defined as an imbalance in calories, macronutrients, vitamins, and minerals during gestation and lactation—can arise from both low-fat, grain-based diets common in resource-limited settings and high-fat, ultra-processed diets. Adequate nutrition is essential for fetal growth, immune development, and postnatal survival.(22-25) For our control diet, we used a custom purified diet that matched the percentage of kcal from fat in the standard chow provided at our institution. There was no statistical difference in septic offspring survival between these two maternal diets; however, survival trended lower using the CD. This supports our use of the custom purified 18% kcal from fat diet as our control diet. Interestingly, the 10%kcal from fat diet is used as a control for the 60% kcal from fat diet by other researchers. Similar to our findings, two studies reported no difference in litter size when dams are fed a 10% or 60% kcal from fat diet.(26, 27) Chen et al. found no difference in P2 offspring born to dams fed a 10% or 60% kcal from fat diet.(26) This difference from our findings, in which offspring of dams fed LFD (10%kcal from fat) weighed significantly less, may be explained by the day of life in which the offspring were weighed. It is possible that the difference in weights we observed is due to differences in milk composition between LFD and CD-fed dams. Despite no differences in maternal weight or body composition between LFD and CD groups, offspring from LFD-fed dams had reduced size and survival at P4-6. It may be that the 10% kcal from fat diet is insufficient to support gestation and lactation. Alternatively, we found that HFD resulted in increased weight and altered body mass composition in dams, with a strong trend toward heavier offspring. This is consistent with some studies but contradicts others, which found lower offspring weight.(28) These variations in both diets may be due to differences in genetic background of mice, time spent on the diet, or variation in diet composition between vendors. Ultimately, offspring of dams fed a LFD or HFD had significantly lower survival from sepsis suggesting that maternal diet fat composition plays an important role in neonatal susceptibility to infection.

Lai et al. found a relatively sublethal dose of 3.5 x10^7 cfu/mL of *S. epidermidis* i.p. produced bacteremia with an inflammatory response in P4 C57BL/6 neonatal mice.(29) They found that serum bacterial counts were significantly elevated at 2h post infection with a mild increase above the detection threshold at 14h and below the detection level by 48h following infection. It is possible that we didn’t find any differences in serum bacterial cfus in septic offspring because of the timepoint we used. Future studies can consider earlier timepoints as later timepoints will be confounded by survival bias as animal deaths start to occur around 12h after infection.

Offspring of both LFD- and HFD-fed dams exhibited reduced basal levels of key cytokines, including IL-6, IL-10, G-CSF, and GM-CSF, indicating a dampened innate immune system prior to infection. Manches et al. did not find any difference in cord blood cytokines among healthy neonates in relation to maternal pre-pregnancy BMI, third-trimester maternal weight, or birth weight.(30) Serum IL-6 is correlated with infection, specifically chorioamnionitis, but not maternal diabetes.(31) Serum IL-10 has been found to be decreased in the serum of children with obesity and rats fed a high-fat diet.(32) Tissue-specific IL-10 levels are also reduced in neonatal mice born to dams fed a high-fat diet.(33) IL-10 is an anti-inflammatory cytokine that helps modulate the inflammatory response. G-CSF and GM-CSF are involved in hematopoiesis, specifically the production and maturation of neutrophils and macrophages. Taken together, this dysregulated basal immune phenotype may predispose neonates to inadequate responses during early infection, which could contribute to the observed decrease in survival.

Neonatal mice, regardless of maternal diet, had significant elevation of a wide array of serum cytokines and chemokines 12 hours following *S. epidermidis* sepsis. Consistent with these findings, Lai et al. found most cytokines and chemokines were significantly increased following *S. epidermidis* sepsis. Nearly all cytokines and chemokines peaked around 14 hours post-infection, except for IL-6 which peaked at 2 hours post-infection, and they returned to baseline by 48h. Interestingly, while both LFD and HFD offspring demonstrated decreased survival, their cytokine responses to sepsis diverged. Offspring of HFD-fed dams exhibited significantly blunted cytokine and chemokine responses during infection, including reduced levels of serum

GM-CSF, TNF-α, IL-1β, IL-6, IL-10, and MCP-1 and peritoneal G-CSF, IL-6, and MCP-1. These findings are consistent with prior studies demonstrating impaired monocyte and dendritic cell responses in neonates born to obese mothers and suggest that maternal overnutrition may induce a state of immune tolerance or exhaustion in the offspring.(16, 34) In contrast, LFD offspring mounted cytokine responses comparable to controls during sepsis. Maternal diet clearly impacted offspring cytokine levels, but there was not a clear relationship between these alterations and sepsis survival, suggesting other mechanisms were involved.

Metabolomic profiling revealed that maternal diet significantly altered the neonatal serum metabolome, with shared significant pathways including phenylalanine and sphingolipid metabolism. These pathways are known to influence immune cell signaling, membrane dynamics, and inflammatory responses, and may represent mechanistic links between maternal diet and offspring immune function.(35-37) Sphingolipid metabolism, specifically the shingosine-1-phosphate metabolite, has been indicated as a potential biomarker and therapeutic target for patients with sepsis.(38-40) The shared metabolic disruptions observed in both LFD and HFD offspring suggest that distinct nutritional insults may converge on common immunometabolic pathways. Future studies will include targeted metabolomic studies to farther explore these pathway differences at a systemic, tissue, and cellular level.

This study has several limitations. First, while we observed significant changes in cytokine levels and metabolite profiles, we did not assess cellular immune responses or tissue-specific inflammation, which may provide further mechanistic insight. Second, our model focused on a single pathogen and time point; future studies should evaluate whether these findings generalize to other infectious challenges and developmental stages. Finally, while our dietary models reflect macronutrient extremes, they do not capture the full complexity of human malnutrition, including micronutrient deficiencies and dietary diversity.

## 5. Conclusion

Our findings highlight the critical role of maternal nutrition in shaping neonatal immune competence and susceptibility to infection. Offspring of dams fed a low-fat or high-fat diet had significantly lower survival from sepsis without a difference in bacterial burden. While offspring of low-fat diet fed dams had a similar cytokine response as offspring of dams fed a control-fat diet, the offspring of high-fat diet fed dams had lower cytokine levels during sepsis, consistent with a dampened immune response. These offspring also had alterations in shared metabolic pathways that may provide a mechanistic reason for their decreased survival during sepsis. These results underscore the importance of optimizing maternal diet during pregnancy and lactation, not only for fetal growth but also for the development of a robust and responsive immune system in the offspring.

## Data Availability Statement

The data that support the findings of this study are available in the Materials and Methods, Results, and/or Supplemental Material of this article.

## Author Contributions

Lauren Bodilly: conceptualization, data acquisition, and data analysis and interpretation. Sarah Weiner: data acquisition. Jennifer Bermick: conceptualization and data analysis and interpretation. All authors were involved in drafting the manuscript and revisions.

This manuscript is the result of funding in whole or in part by the National Institutes of Health (NIH). It is subject to the NIH Public Access Policy. Through acceptance of this federal funding, NIH has been given a right to make this manuscript publicly available in PubMed Central upon the Official Date of Publication, as defined by NIH.

